# Visual and oculomotor abilities predict professional baseball batting performance

**DOI:** 10.1101/2020.01.21.913152

**Authors:** Sicong Liu, Frederick R. Edmunds, Kyle Burris, Lawrence Gregory Appelbaum

## Abstract

Scientists and practitioners have long debated about the specific visual skills needed to excel at hitting a pitched baseball. This study aimed to advance the debate by evaluating the relationship between pre-season visual and oculomotor evaluations and pitch-by-pitch season performance data from professional baseball batters. Eye tracking, visual-motor, and optometric evaluations collected during spring training 2018 were obtained from 71 professional baseball players. Pitch-level data from Trackman 3D Doppler radar were obtained from these players during the subsequent season and used to generate batting propensity scores for swinging at pitches out of the strike zone (O-Swing), swinging at pitches in the strike zone (Z-Swing), and swinging at, but missing pitches in the strike zone (Z-Miss). Nested regression models were used to test which vision-related evaluation(s) could best predict the standardized plate discipline scores as well as the batters’ highest attained league levels during the season. Results indicated that visual evaluations relying on eye tracking (e.g., smooth pursuit accuracy and oculomotor processing speed) significantly predicted the highest attained league level and the propensity scores associated with O-Swing and Z-Swing, but not Z-Miss. These exploratory findings indicate that batters with superior visual and oculomotor abilities are generally more discerning at the plate. When combined with other known performance advantages in perceptual and cognitive abilities for elite athletes, these results provide a wholistic view of visual expertise in athletes.

## 1. Introduction

Hitting a pitched ball is among the most iconic of all sporting activities. Across baseball, softball, cricket and other batting sports, the rules of the game have created situations that very precisely test the limits of human’s abilities to see and react. By specifying the dimensions of the strike zone, and the distance and height between the pitcher and batter, these sports have created scenarios where the competitive balance of the game unfolds over a few hundred milliseconds, with balls moving at peak velocities exceeding the capacity of the human oculomotor system (Spering & Gegenfurtner, 2008; Watts & Bahill, 1991). The extreme challenge of this endeavor is punctuated by the fact that hitting successfully in one out of three plate appearances can garner contracts that now routinely exceed twenty million dollars a year.

In baseball, for example, fastball pitches regularly exceed 95 miles per hour, traveling approximately 55 feet from the pitcher’s hand to home plate in under 350 milliseconds. Through this process, the batter must decipher the pitch, project its trajectory, decide to swing or not, and coordinate the timing and movement of a 2.25-inch diameter bat to intercept a 3-inch ball. To have the best chance to hit the ball, batters look for cues that tip the pitch during the wind up, like the placement of the pitchers fingers relative to the seams, and extract movement information from the arm and the ball, including the spin, to project the trajectory relative to the strike zone.

The unique skills that allow expert batters to accomplish these feats has been an area of substantial scientific interest. Studies contrasting batters at different achievement levels show that experts demonstrate better pitch anticipation than non-experts, and that such pitch anticipation positively correlates to batting statistics (Müller & Fadde, 2016). It has been shown that anticipating the pitch type at the moment the ball is released relies on reading pitcher kinematics, particularly those in the hand-shoulder region (Kato & Fukuda, 2002), and that anticipating where the pitch eventually crosses the plate entails tracking ball flight early in its path, at least over the first 80 milliseconds (Müller et al., 2017; Paull & Glencross, 1997). Moreover, studies involving eye tracking revealed that expert batters utilize information from early ball tracking to generate predictive saccades to place the eye ahead of the trajectory of the moving ball (Bahill & Laritz, 1984; Land & McLeod, 2000; D. L. Mann et al., 2013). In this way the batter can ‘wait’ for the ball to enter the visual field, circumventing the problem of tracking a ball that moves faster than the oculomotor system can resolve (Spering & Gegenfurtner, 2008).

Collectively these abilities reflect a combination of receptive visual abilities that transform the light signal into the neural code and perceptual visual abilities that process the input for meaning, context, intention, and action, so-called visual “hardware” and “software”. While there is considerable evidence that software abilities such as anticipation, pattern recognition, and visual search, are elevated in higher performing athletes (see meta-analyses by Lebeau et al., 2016; D. Y. Mann et al., 2007; Voss et al., 2010), there is less evidence linking visual-hardware to greater athletic expertise. Therefore, while there is evidence that visual acuity (Laby et al., 1996) and contrast sensitivity (Hoffman et al., 1984), are better in higher-level athletes, there is still an incomplete picture of how these traits might impact performance.

Given the potential value of establishing characteristics that predict future performance in baseball, there has been a growing effort to map specific visual skill to on-field batting performance. For instance, visual-motor skills tested on the Nike Sensory Station were shown to predict several game statistics including on-base percentage, walk rate, and strikeout rate (Burris et al., 2018). Additionally, athletes with better dynamic visual abilities, those that rely on acuity and contrast sensitivity judgments performed under temporal constraints, were shown to produce better ‘plate discipline’ batting statistics (e.g., O-Swing Propensity, which is discussed in the Methods section below; Laby et al., 2019). Furthermore, when compared between batters and pitchers with similar levels of experience, batters were shown to produce better performance on measures of visual acuity and depth perception than pitchers (Klemish et al., 2018), indicating that these skills are specific to the demand of hitting pitched ball, not throwing them. Lastly, eye tracking research suggested that batters shifting visual fixations more frequently between pitcher and home plate prior to batting showed better on-base percentage and batting average (Hunfalvay et al., 2019).

While the studies presented above have provided novel and systematic insight into the relationship between visual abilities and batting performance, they have tended to use relatively narrowly-defined visual and/or perceptual-cognitive assessments, resulting in dependence on specific assessment modality (e.g., responding with hand-held devices). In addition, they typically focused on batters’ performance measures (e.g., on-base percentage) that do not control for the contribution of the defense (though see Laby et al., 2018). In the present study, we aimed to improve upon these limitations by making use of a wide range of assessments, collected as part of pre-season evaluations that include measures of refractive error, quantitative eye tracking, and visual-motor abilities. Such an array of visual assessments allows for comparison of the relative importance among assessment modalities, in addition to the visual constructs that are captured by the test batteries. To infer the role of these abilities on batting performance, we make use of pitch-by-pitch plate discipline metrics (collected during the subsequent season) that rely only on the batter’s abilities and are not influenced by the fielder’s defensive performance. By mapping a broad range of visual assessments to context-controlled plate discipline statistics this study aims to clarify which aspect of visual skill, in which assessment modality, contribute the most to batting performance. It was hypothesized that superior visual assessments would correspond to better baseball performance.

## 2. Methods

### 2.1. Participants

The study included a sample of 71 professional minor league baseball batters (*M* = 22.1 years, *SD* = 2.5 years). **Table 1** reports the sample distribution regarding handedness and League Level, defined as the highest minor league level attained by a given batter during the 2018 season. Multiple empirical datasets contributed to obtaining the sample and they formed two general categories, one including visual assessment (see 2.2. Visual Assessments) and the other involving pitch-by-pitch performance measured throughout the 2018 season (see 2.3. Plate Discipline Variables). Merging the datasets via encrypted IDs led to an initial sample of 109 batters. This dataset was further adjusted because of missing values and potential collinearity issues among visual assessment variables. Specifically, only batters with less than 10 missed observations out of a total of 22 visual assessment variables were included, and the number of visual assessment variables were reduced to 14 by generating composite variables among those of high bivariate correlation (i.e., Pearson *r*s > .50) and conceptual relatedness (see Section 2.2). These steps resulted in a final sample of 71 batters and an overall missing data rate of 3.1% in the final dataset.

**Table 1:**
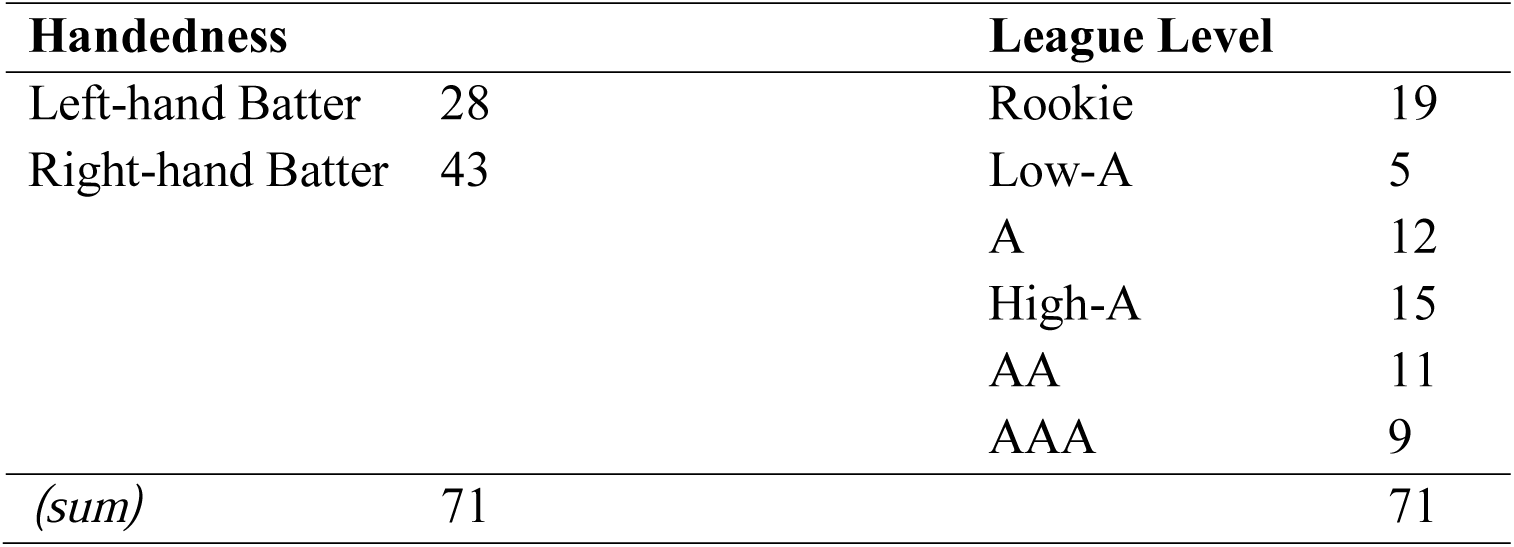
Sample characteristics of included players.

All data were shared under a secondary-data protocol [IRB B0706] approved by the Duke University Institutional Review Board and the Human Research Protection Office of the US Army Medical Research and Materiel Command under separately reviewed protocol. Under these protocols, all data were collected for “real world use,” without informed consent, and shared via encrypted IDs without inclusion of any protected health information. These data therefore conform to U.S. Department of Health and Human Services, “Regulatory considerations regarding classification of projects involving real world data”(DHHS, 2015) and also to the ethical principles of the Declaration of Helsinki.

### 2.2. Visual Assessments

Assessments were performed between March and May 2018, by an optometrist retained by the professional baseball franchise (author FE). Assessments took place at the team’s spring training facility, primarily during the player’s first two weeks of training camp. Three separate evaluation stations were set up to capture visual assessment data. The assessments at each station took between 5-12 minutes to complete and while these were generally done sequentially, occasionally, a player’s assessment took place over the course of two days if they were unable to complete all stations in one day. Instructions for each assessment were given in English and when necessary were augmented with instructions in Spanish. In each case, demonstrations and practice were given with the equipment prior to testing.

#### 2.2.1. Eye-Tracking Assessments

Quantitative eye tracking was performed using customized sub-sets of the RightEye LLC (Bethesda, MD), Neuro and Performance Vision assessment batteries. Testing was performed in a private, quiet testing room with participants seated with their eyes in front of a NVIDIA 24-inch 3D Vision monitor at a distance of 60 cm. Eye tracking was achieved through a 12-inch SMI, 120 Hz remote eye tracker, connected to an Alienware gaming system and a Logitech (model Y-R0017) wireless keyboard and mouse. Participants heads were unconstrained during the test, although they were instructed to sit still and warned with indicators on the screen if the eye tracking was interrupted. The RightEye test battery began with a digital confirmation that the eyes were centered on the screen (with a feedback icon to facilitate adjustment), followed by a nine-point calibration test in which tracking fidelity was evaluated over the full expanse of the screen. Upon successful calibration, the task battery commenced. For each successive task, text and animated instructions were provided. Detailed information about the tasks can be found elsewhere (Murray et al., 2019) and the following measures were calculated from performance on the test battery for analysis in the current study:

- **Dynamic Visual Acuity** is a measure of the ability to recognize fine details of an object moving across a monitor screen while the participant is instructed to keep their head still. Performance is quantified in seconds, with smaller values reflecting better performance.
- **Cardinal Reaction Time** is the time required for participants to move visual gaze from the central fixation mark to pictorial targets appearing at eight cardinal positions. Performance is quantified in seconds, with smaller values indicating better performance.
- **Simple Reaction Time** is the time required to press a keyboard button in response to presentation of pictorial targets displayed at fixation. Responses are measured in seconds with smaller values indicating better performance.
- **Smooth Pursuit Accuracy** is a composite variable reporting the percentage of time that participants are able to maintain their gaze within three degrees of a smoothly moving black target dot. This measure is calculated as the average over three pursuit trajectories; circular, horizontal and vertical with larger values indicating better performance.
- **General Oculomotor Latency, General Oculomotor Speed**, and **General Processing Speed** are three composite time variables calculated by averaging similar measures from a choice reaction time task and a discriminant reaction time task. Both tasks require participants to fixate centrally, locate and foveate on incoming targets that project inward from the periphery of the screen, and make manual responses to indicate the identity of the target once acquired visually. Oculomotor Latency refers to the elapsed time from the appearance of the target to the moment gaze is averted from the central fixation mark. Oculomotor Speed refers to the elapsed time from the moment when gaze is averted from the central fixation to the moment when gaze arrives at the incoming pictorial target. Processing Speed refers to time elapsed between arrival of gaze to the incoming target and the moment when responses are registered with a keyboard button press. Measures are referred to as “general” because they are averaged over identical metrics for two tasks, in order to create a more robust measure that captures this construct. All are measured in seconds with smaller values indicating better performance.

#### 2.2.2. Visual-Motor Assessments

Visual-motor skills were assessed using the Senaptec LLC (Beaverton, OR) Sensory Station Tablet. This device presents a battery of computerized visual-motor tasks, each designed to evaluate a specific facet of a participant’s visual-motor abilities. Testing was performed at two distances; 10 feet and 18”-24”. The tablet was mounted on a sturdy, adjustable tripod, with the center of the screen positioned at eye level. Tasks performed at distance were conducted by the participants using a remote controller connected to the Tablet via Bluetooth. The remaining tasks were performed by the participant directly on the tablet touch screen at arm’s length. Detailed information about the tasks can be found elsewhere (Wang et al., 2015) and the following measures were calculated from performance on the test battery for analysis in the current study.

- **Visual Clarity** is a measure of static visual acuity obtained by having participants report the orientation of gaps in a Landolt ring, presented at distance, and adjusted in size according to an adaptive staircase procedure. Scores are reported in LogMAR units with smaller values indicating better performance.
- **Contrast Sensitivity** measures the minimal lightness-darkness contrast shown in static ring-shaped targets displayed at distance. Stimuli are presented at 18 cycles-per-degree and adjusted in contrast according to an adaptive staircase procedure. Results are reported in log contrast with larger values indicating better performance.
- **Near-Far Quickness** is a measure of how quickly participants could visually accommodate back and forth between near and far visual targets in 30 seconds, without
- sacrificing response accuracy. Scores indicate the number of correctly reported targets with larger values indicating better performance.
- **Multiple Object Tracking** is a measure of how well participants could maintain accurate spatial tracking of multiple moving targets, presented with moving non-targets according to an adaptive staircase schedule. Scores are computed as a composite of movement speed thresholds and tracking capacity, with larger values indicating better performance.
- **Perception Span** is a measure of spatial working memory derived by having participants recreate the locations of briefly presented targets that are flashed in a grid of circles. The number of targets and the size of the grid increases with correct responses, and the final score indicates the combined total of correct responses, minus errors, across all levels. Larger values indicate better performance.
- **Reaction Time** is the elapsed time between when rings on the touch pad change color, and when participants are able to remove their index finger from the touch sensitive screen. Scores are reported in seconds with smaller values indicating better performance.

#### 2.2.3. Auto-Refraction

An Ovitz (West Henrietta, NY) P10 autorefractor was utilized to capture objective measurement of the refractive error of each eye, for each individual. Measurements were taken in a dimly-lit and quiet room and participants were instructed to fixate on a target placed 10 feet away to control and minimize accommodation. The Ovitz device was alternately positioned in front of the right eye (R), then left eye (L), allowing participants to maintain a far focus with the non-fixating eye. Using the individual left and right eye spherical (sph) and cylindrical (cyl) values, spherical equivalence was calculated using the following formula:

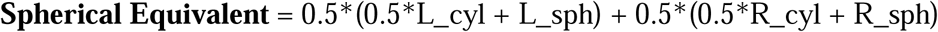

### 2.3. Plate Discipline Variables

Batters’ pitch-by-pitch data based on 3D Doppler radar systems (Trackman LLC., Stamford, CT) are valuable in quantifying on-field performance because they reflect the precise trajectory of each pitched and hit baseball. In this study, batters’ pitch-by-pitch data was linked to their vision assessments by the franchise’s analytics department and shared with the research team in a deidentified manner. These data were then used to model batters’ propensity scores for three plate discipline variables including:

- **O-Swing %**, defined as the number of swings at pitches outside the strike zone divided by the number of pitches seen outside the strike zone. Lower values are preferred.
- **Z-Swing %**, defined as the number of swings at pitches inside the strike zone divided by the number of pitches seen inside the strike zone. Lower values indicate more discerning batters.
- **Z-Miss %**, defined as the number of missed swings at pitches inside the strike zone divided by the number of swings at pitches inside the strike zone. Lower values are preferred.

The modeled propensity scores can be viewed as a standardization of the above plate discipline variables that accounts for player heterogeneity in the difficulty of pitches faced. For example, a player in AAA is more likely to face pitches of near MLB quality, resulting in far more swings and misses than at the Rookie level. By accounting for game context in this way, the propensity scores tend to be more accurate representations of underlying player ability than the raw percentages (Gray, 2002b). Therefore, in order to isolate a batter’s batting ability, we control for the quality and context of pitches faced via a generalized additive mixed model (GAMM; Wood, 2004). To obtain estimates of each player’s propensity score on a given plate discipline variable, a GAMM was fit to all the available pitches with player-specific random effects. The model also included cubic spline terms and tensor interactions to account for the location of the pitch, the count, the movement of the pitch, the speed of the pitch, the spin rate of the pitch, the handedness of the batter, and the handedness of the pitcher. The response variable was a binary indicator for either a swing decision or the batter making contact with the ball, depending on the statistic-of-interest. For instance, the GAMM for swing-chase propensity was trained on all pitches thrown outside the strike zone and models the probability of a swing. The estimated values of the random effects for the players were extracted and treated as the propensity score for the underlying statistic. In general, these propensity scores were nearly normally distributed across players in the sample (see bottom row of **Figure 1**).

**Figure 1.**
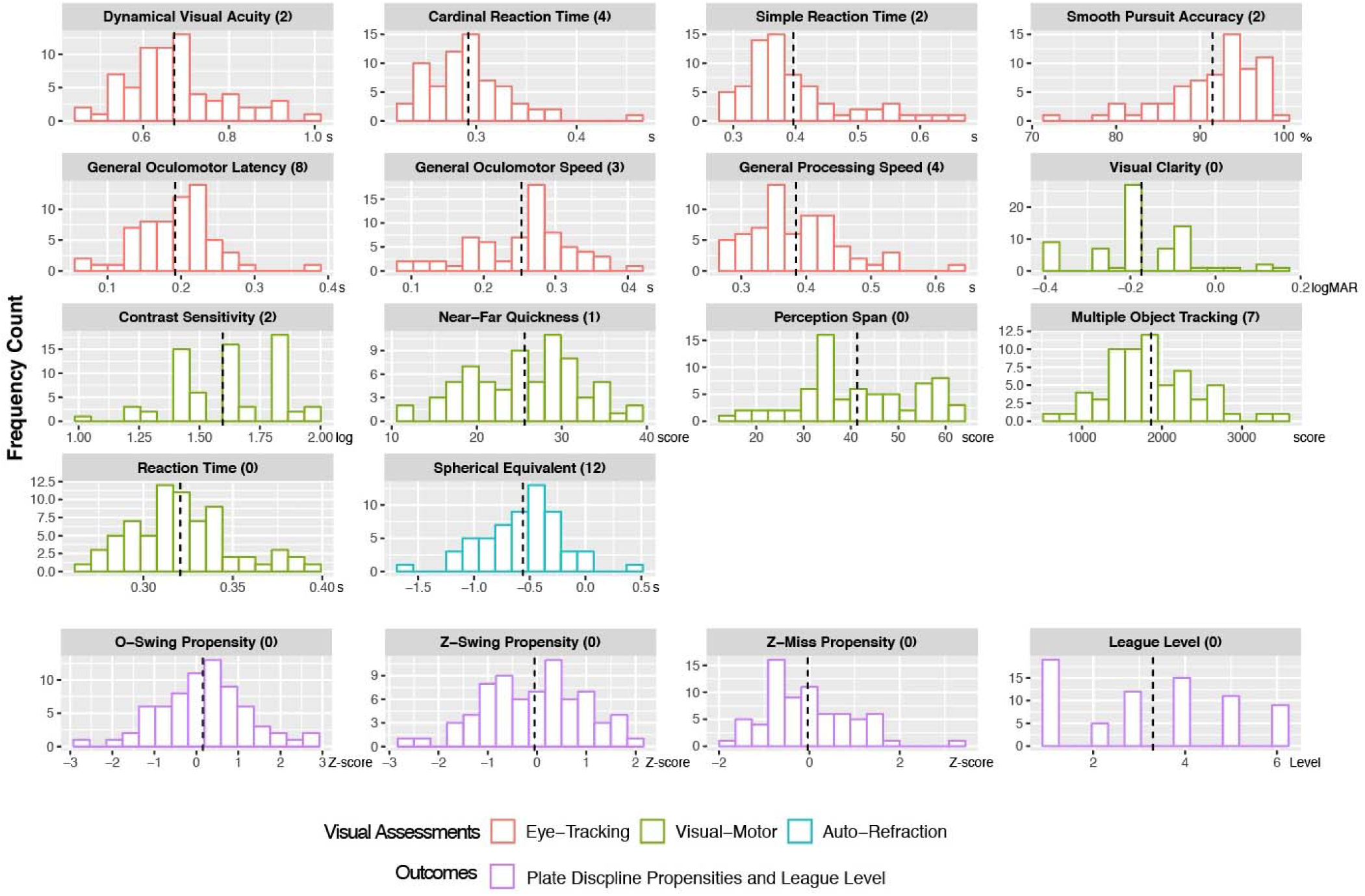
Variable histograms with mean values indicated by the dashed lines and the number of missing values (out of 71) in parentheses. Units are indicated at the bottom right of each X-axis labels.

### 2.4. Analytical Approach

All analyses were conducted using R 3.5.3 (R Core Team, 2019). A preliminary analysis was first performed to check for violations of statistical assumptions. Seven suspected outliers from five visual assessment variables (i.e., Contrast Sensitivity, Near-Far Quickness, Cardinal Reaction Time, Simple Reaction Time, General Processing Speed) were removed. The outlier judgements were informed by the lower and upper whisker length of 1.5 inter-quarter range (IQR) in the boxplot. Removing outliers increased the overall missing rate to 3.7% in the empirical dataset prior to multiple imputation. **Figure 1** displays the histogram of all the variables used in the nested regression models prior to multiple imputation. Multiple imputation procedures were implemented using the *Mice* package in R. Specifically, 20 imputed datasets were generated with the method of *Predictive Mean Matching* and a maximum of 50 iterations.

Given the exploratory nature of the study, a set of nested regression models was planned for each plate discipline variable. That is, the propensity scores for each plate discipline variable were regressed on the visual assessment variables in a nested structure, based on the evaluation modality of the vision-related variables. The nested models are listed below:

- Model 1: Plate Discipline ∼ (intercept) + Eye-Tracking + Visual-Motor + Auto Refraction
- Model 2: Plate Discipline ∼ (intercept) + Eye-Tracking + Visual-Motor
- Model 3: Plate Discipline ∼ (intercept) + Eye-Tracking
- Null Model: Plate Discipline ∼ (intercept)

To validate the predictive power of visual assessment variables on another measure of baseball success, a similar set of nested regression models was also performed on the League Level variable. Because League Level is ordinal, ordinal logistic regression was chosen instead of multiple regression for this analysis. All the nested regression models were fit to each of the 20 imputed datasets. To assure that the sequential nesting of variables in the model did not mask effects of visual-motor assessments, an additional model was tested: Plate Discipline ∼ (intercept) + Visual-Motor. However, such a model showed descriptively smaller *R*^2^ values than Model 3 and no statistically better fit than the Null Model on all the outcome variables (i.e., propensity scores associated with plate discipline variables and league level). The model was thus dropped from further considerations.

For a given regression model, its final parameter estimates were based on pooling the estimates generated when fit to the imputed datasets (D. B. Rubin, 1987). This pooling helped account for the uncertainty associated with the missing data values. The nested models for a given outcome were treated as competing ones and compared based on overall goodness-of-fit. In particular, Wald’s test was used to compare nested models on a given plate discipline propensity and likelihood ratio X^2^ test was used for comparing nested models on League Level. For a given set of nested model test, a final model was considered when showing reasonable overall goodness-of-fit, accounting for meaningful variance in the outcome, and remaining parsimoniously specified (i.e., minimizing the number of parameters in a given model). Although the alpha level was set at .05, model comparisons whose *p-values* approached this level (i.e., *p* < .10) were considered in presence of other favorable evidence (e.g., effect size and parsimoniousness).

## 3. Results

Because four nested models were specified for a given outcome variable, model comparisons and selections were necessary to reach the final model that balanced parsimony and explanative power. **Table 2** shows the *p*-*values* associated with the model comparison tests and the *R*^*2*^ estimate for each given model. Both these results were taken into consideration when selecting the final model for each outcome variable. **Figure 2** illustrates the statistical significance of individual visual assessment variables in each selected regression model for a given outcome variable. The Supplemental Contents (SCs) contain model fits for the individual visual assessment variables (SC1), confidence intervals for the *R*^*2*^ estimates (SC2), and the *p* values from comparing vision models on a given outcome variable (SC3).

**Table 2:** p-values associated with vision-vs-null model comparisons and R^2^ estimates for the vision models

| | *P* value | | | $R^2$ value | | |
| --- | --- | --- | --- | --- | --- | --- |
|  | Model 1<br>vs. Null | Model 2<br>vs. Null | Model 3<br>vs. Null | Model 1 | Model 2 | Model 3 |
| O-Swing Propensity | 0.56 | 0.38 | 0.07 | 0.33 | 0.33 | 0.28 |
| Z-Swing Propensity | 0.07 | **0.05** | 0.35 | 0.23 | 0.22 | 0.09 |
| Z-Miss Propensity | 0.29 | 0.3 | 0.95 | 0.21 | 0.19 | 0.03 |
| League Level | < .001 | < .001 | < .001 | 0.16<sup>†</sup> | 0.15<sup>†</sup> | 0.13<sup>†</sup> |

Note that text is bolded if *p*-values were less than .05, and boxed cells indicate models that were further evaluated. <sup>†</sup> pseudo- $R^2$ s ( ) values, calculated using model deviance estimates based on measures of likelihood, whose interpretation is not the same as $R^2$ and requires caution.

**Figure 2.**
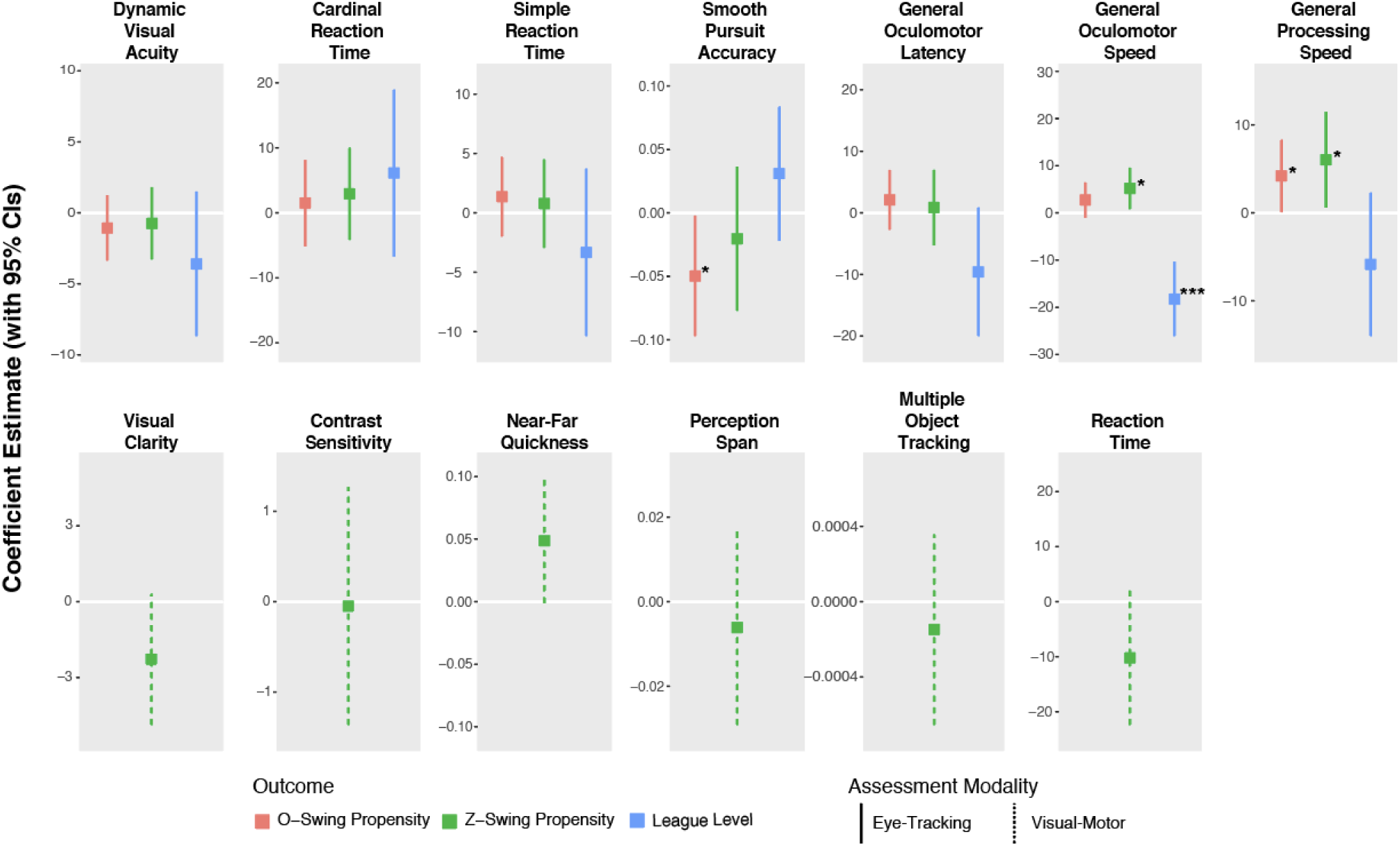
Slope parameter estimates and 95% confidence intervals (CIs) of visual assessment variables in the selected regression models on O-Swing Propensity (red), Z-Swing Propensity (green), and League Level (blue). Gen. = General. * p < .05, *** p < .001.

### O-Swing Propensity

Model 3 was selected for further consideration because it was the only tested model showing a marginally significantly better fit than the Null Model (*p* = .07), while demonstrating substantial predictive power wherein the visual assessment variables accounted for nearly 28% of the variance in chase rates, *R*^2^ = .28. Model 3 results revealed that eye-tracking variables of Smooth Pursuit Accuracy, *t*(56.84) = -2.10, *p* = .04, and General Processing Speed, *t*(56.58) = 2.04, *p* = .046, were significant predictors for O-Swing Propensity. This observation indicates that better smooth pursuit accuracy and faster information processing speed were associated with lower propensity to swing at pitches outside the strike zone.

### Z-Swing Propensity

Model 2 was selected for further consideration as it showed a significantly better fit than the Null Model, *p* = .05, while demonstrating substantial predictive power from visual assessment variables, accounting for 22% of the variance, *R*^*2*^ = .22. Model 2 results indicated that eye-tracking variables of General Oculomotor Speed, *t*(53.64) = 2.37, *p* = .03, and General Processing Speed, *t*(53.49) = 2.23, *p* = .03, were significant predictors for Z-Swing Propensity. In both cases, better performance on these assessments corresponded to lower swing rates for pitches inside the strike zone, implying that individuals with better visual abilities are more discerning in their swings and tend to swing less at pitches.

### Z-Miss Propensity

None of the tested models showed better fit than the Null Model and thus no final model was selected for this plate discipline variable.

### League Level

Model 3 was selected for further consideration due to its parsimony compared to Model 1 and Model 2, while all the three tested models demonstrated better fits than the Null Model, *p*s < .001. Model 3 results demonstrate that the eye-tracking variable of General Oculomotor Speed, *t*(53.48) = -4.66, *p* < .001, was a significant predictor of League Level, while General Oculomotor Latency, *t*(43.83) = -1.85, *p* = .07, trended towards significance. These findings indicated that athletes who reach higher leagues tend to have faster oculomotor movement speeds.

## Discussion

Phrases like “keep your eyes on the ball” and “you can’t hit what you can’t see” underscore the important role that visual perception plays in baseball performance. Despite contributions from past studies, uncertainty still exists about the nature of visual skills that contribute to hitting performance in such highly demanding interceptive actions. In particular, while several studies have linked better anticipatory skills with greater batting performance (Kato & Fukuda, 2002; Müller et al., 2017; Müller & Fadde, 2016; Paull & Glencross, 1997), less is known about the role of oculomotor and visual-perceptual skills. The current study aimed to contribute to this area of understanding by evaluating the links between visual skills and plate discipline statistics among professional baseball batters in a naturalistic dataset. Validated visual assessments based on auto-refraction (A. Rubin & Harris, 1995), eye-tracking (Murray et al., 2019), and visual-motor control (Wang et al., 2015), commissioned by a professional baseball franchise during the preseason, were mapped to plate discipline performance modeled using Trackman pitch-by-pitch data collected throughout the ensuing season. Due to the exploratory nature of the study, several competing models and the null model were tested against each other on criteria of overall goodness-of-fit, model parsimoniousness, and effect size (i.e., *R*^*2*^).

Results demonstrate that, compared to auto-refraction and visual-motor measures, several oculomotor eye tracking measures stood out in predicting plate discipline performance and the athlete’s highest attained league level. In particular, batters with faster oculomotor and processing speeds, as well as better smooth pursuit accuracy, tended to be more discerning in their swings by lowering the swing propensity regardless of pitch location (i.e., inside/outside strike zone), while batters with faster oculomotor speeds also tended to compete in higher professional leagues. Given the observation that batters who are less likely to swing at strikes actually show a higher chance of making ball contact to generate fair plays and obtain walks (Albert, 2017), this pattern of findings is consistent with the hypothesis that better visual skills predict better plate discipline performance (with the exception for Z-Miss Propensity), as well as higher achieved League Level.

The present evidence informs practitioners through the comparison of a relatively wide range of visual assessments that can be linked to performance in baseball batting. Specifically, visual measures based on eye tracking emerged with predictive potential on both O-Swing Propensity (*R*^*2*^ = 0.28) and Z-Swing Propensity (*R*^*2*^ = 0.22). The final model implies that an individual with 1% better smooth pursuit accuracy, or 10 ms faster oculomotor/processing speeds would be 2% more discerning in swinging relative to his peers. Oculomotor speed also predicted the highest attainted league level, a different means to characterize baseball proficiency. According to the estimated slope parameter of the odds ratio metric in the final ordinal logistic regression model, a batter will be 1.22 times more likely to play at a higher league levels than lower ones if the batter has 10 ms faster oculomotor speeds. An important next step in this exploratory research will be to validate the current findings in independent samples. Contingent on such replications, these findings have important implication for the promise of utilizing oculomotor assessments as a means to scout baseball batters based on these evaluations.

The finding that oculomotor and information processing speeds, as well as smooth pursuit accuracy, were most predictive on O-Swing and Z-Swing Propensity bears several conceptual implications. First, as previously outlined, expert batters need to maintain ball tracking for at least 80 ms after the moment of pitch release to anticipate the pitch location at above-chance levels (Müller et al., 2017). Faster information processing speeds and enhanced pitch tracking accuracy, especially at the start of the pitch flight while the ball is traveling at the lowest range of angular velocity, may help batters lower O-Swing Propensity through sharpened recognition of pitches outside the strike zone. Similarly, the sharpened pitch-location recognition through faster oculomotor and information processing speeds may also help batters be more discerning when facing pitches inside the strike zone. In a follow-up analysis of the current sample, we obtained a marginally significant positive bivariate correlation (*r* = .18, *p* = .06) between Z-Swing Propensity and Z-Miss Propensity, implying that a more discerning batter is also more likely to make contact with pitches going inside the strike zone. This view is consistent with the idea that swing actions from more discerning batters are more likely to result in hitting into fair plays (Laby et al., 2019).

Second, better recognition of pitch location is probably not sufficient to make a batter more discerning in swinging, given that elite batters tend to “sit on fastballs” (Gray, 2002b). Sitting on a fastball refers to a proactive strategy of baseball batters to always anticipate a fastball because they are most common and require the greatest speed challenge (Canãl-Bruland et al., 2015), creating a bias towards swinging. Considering that faster processing speed characterizes greater competence in switching responses according to visual stimuli, batters with faster information processing speed may be more discerning in their swings because they can better inactivate the swing initializations triggered by “sitting on fastballs” (Muraskin et al., 2015).

Third, the modest *R*^2^ values obtained in the most complex model group (see those of Model 1 in **Table 2**) indicate that factors other than the included visual assessments are contributing substantially to the plate discipline measures. For instance, baseball batters are reported to utilize information from auditory and tactile feedback in performance, although they rely more on visual feedback (Gray, 2009). Within the current range of visual assessments, information redundancy also exists, consistent with past studies reporting that the majority of variance in a battery of nine visual-motor tasks developed by Nike is accounted for by three latent factors (Poltavski & Biberdorf, 2015; Wang et al., 2015). In the current study, for instance, despite the fact that many baseball teams work with optometric professionals, the clinical measure of auto-refraction was not found to predict plate discipline measures or highest attained league level given the presence of other visual assessments in the models. It is possible that good optometric assessments are only necessary conditions for batting excellence, but, among batters who demonstrate superior vision as a group, the utility of auto-refraction to capture variance in batting performance might have been limited.

Lastly, none of the assessed visual skills were found to predict Z-Miss Propensity. This result is interesting because it may suggest strong determinants of swinging and missing, other than visual skills. Since it has been reported that striking a 90 mile/hour fastball requires keeping the temporal error within ± 9 ms and spatial error within ± 1.27 cm (Gray, 2002a; Regan, 1997), it could be that Z-Miss propensity relies on not only ‘seeing’ the pitch but also swinging with temporal and spatial accuracy (Lee, 1998). Therefore, the ability of making accurate swing actions may be a test-worthy factor together with visual skills for predicting Z-Miss Propensity in future.

### Future Directions and Limitations

The present findings must be taken within the context of several limitations, but also invite a number of important future research questions. First, as noted above, the analytical approach was exploratory wherein alternative models were considered in the presence of multiple sources of evidence, including goodness-of-fit, parsimony, and effect size. The study included a ‘convenience sample’ enabled through a research collaboration, and therefore there is a strong need to replicate the present findings in independent, larger and experimentally-controlled samples. With the recent adoption of eye tracking measures by USA Baseball (RightEye, 2019) in the player development pipeline, there should be potential opportunities for replication and further investigation using the current exploratory findings as the basis for more explicit hypothesis tests.

Second, although the current study included a wide range of visual skill assessments, other important abilities may have been missed in these evaluations. For instance, parafoveal visual skills have been shown to vary across individuals with some individuals demonstrating a strategy of “parafoveal tracking” when facing pitches in cricket (Croft et al., 2010) and similar strategies in other interceptive sports, such as table tennis (Ripoll & Fleurance, 1988). Therefore, it may be important to include parafoveal visual skills in future assessment battery.

Third, the current findings provide a useful framework for understanding which visual skills are important for batting performance, opening the door for innovations in vision-based and/or virtual-reality-based training protocols to improve batting performance. In particular, there has been rapid development of digital training tools that are based on perceptual learning protocols that can deployed in natural training contexts to promote sports-specific visual and cognitive abilities (Appelbaum & Erickson, 2018; Wilkins & Appelbaum, 2019). With the increased use of these training programs, however, it will be important for research to adhere to a greater level of scientific rigor, including pre-registration, randomization, and placebo control, all features of a current study underway by our research team (Appelbaum et al., 2018). Nonetheless, the finding of present study may serve as a stepping stone for future training studies by highlighting visual skills that can be targeted in such interventions.

Lastly, while the current findings point towards possible use of oculomotor assessments as a means to scout batting talent, there are likely causal factors that influence the development of these skills not testable in the current design. As such, it remains an open question as to whether batters reach higher league levels because natural abilities, or if these capabilities are honed over an athlete’s development. Future longitudinal studies may help to arbitrate this question.

## Conclusion

The present exploratory research findings indicate that oculomotor skills predict specific baseball hitting abilities. These findings suggest a possibly valuable source of scouting data and targets for vision-based training programs that may improve batting performance. As, such, future hypothesis-driven research may use the characteristics identified here to guide studies testing talent identification or training studies aimed at improving on-field performance.

## Supporting information

Supplemental Contents

## Acknowledgments

The authors would like to thank the players, coaches, trainers and management who contributed to this study.

## References

Albert, J. (2017). Visualizing baseball. In Visualizing Baseball. https://doi.org/10.1201/9781315149530

Appelbaum, L. G., & Erickson, G. (2018). Sports vision training: A review of the state-of-the-art in digital training techniques. International Review of Sport and Exercise Psychology, 11(1), 160–189. https://doi.org/10.1080/1750984X.2016.1266376

Appelbaum, L. G., Liu, S., Hilbig, S., Rankin, K., Naclario, M., Asamoa, E., LaRue, J., & Burris, K. (2018). Sports vision training in collegiate baseball batters. https://doi.org/10.17605/OSF.IO/496RX

Bahill, A., & Laritz, T. (1984). Why can’t batters keep their eyes on the ball? American Scientist, 72(3), 249–253. https://doi.org/10.1186/scrt73

Burris, K., Vittetoe, K., Ramger, B., Suresh, S., Tokdar, S. T., Reiter, J. P., & Appelbaum, L. G. (2018). Sensorimotor abilities predict on-field performance in professional baseball. Scientific Reports, 8(1), 116. https://doi.org/10.1038/s41598-017-18565-7

Canãl-Bruland, R., Filius, M. A., & Oudejans, R. R. D. (2015). Sitting on a fastball. Journal of Motor Behavior, 47(4), 267–270. https://doi.org/10.1080/00222895.2014.976167

Croft, J. L., Button, C., & Dicks, M. (2010). Visual strategies of sub-elite cricket batsmen in response to different ball velocities. Human Movement Science, 29(5), 751–763. https://doi.org/10.1016/j.humov.2009.10.004

DHHS. (2015). Secretary’s advisory committee on human research protections. Attachment A: Human subjects research implications of “big data” studies.

Gray, R. (2002a). Behavior of college baseball players in a virtual batting task. Journal of Experimental Psychology: Human Perception and Performance, 28(5), 1131–1148. https://doi.org/10.1037/0096-1523.28.5.1131

Gray, R. (2002b). “Markov at the bat”: A model of cognitive processing in baseball batters. Psychological Science, 13(6), 542–547. https://doi.org/10.1111/1467-9280.00495

Gray, R. (2009). How do batters use visual, auditory, and tactile information about the success of a baseball swing? Research Quarterly for Exercise and Sport, 80(3), 491–501.

Hoffman, L. G., Polan, G., & Powell, J. (1984). The relationship of contrast sensitivity functions to sports vision. Journal of the American Optometric Association, 55(10), 747–752.

Hunfalvay, M., Roberts, C.-M., Ryan, W., Murray, N., Tabano, J., & Martin, C. (2019). An investigation of the visual behavior of professional baseballers prior to the execution of batting: Evidence for oculomotor warm up, attentional processes or pre-performance routines? International Journal of Sports Science, 7(6), 215–222. https://doi.org/10.5923/j.sports.20170706.02

Kato, T., & Fukuda, T. (2002). Visual search strategies of baseball batters: Eye movements during the preparatory phase of batting. Perceptual and Motor Skills, 94(2), 380–386.

Klemish, D., Ramger, B., Vittetoe, K., Reiter, J. P., Tokdar, S. T., & Appelbaum, L. G. (2018). Visual abilities distinguish pitchers from hitters in professional baseball. Journal of Sports Sciences, 36(2), 171–179. https://doi.org/10.1080/02640414.2017.1288296

Laby, D. M., Kirschen, D. G., & Govindarajulu, U. (2019). The effect of visual function on the batting performance of professional baseball players. In Press. Scientific Reports, 9, 16847.

Laby, D. M., Kirschen, D. G., Govindarajulu, U., & Deland, P. (2018). The hand-eye coordination of professional baseball players: The relationship to batting. Optometry and Vision Science, 122(4), 476–485. https://doi.org/10.1097/OPX.0000000000001239

Laby, D. M., Rosenbaum, A. L., Kirschen, D. G., Davidson, J. L., Rosenbaum, L. J., Strasser, C., & Mellman, M. F. (1996). The visual function of professional baseball players. American Journal of Ophthalmology, 122(4), 476–485. https://doi.org/10.1016/S0002-9394(14)72106-3

Land, M. F., & McLeod, P. (2000). From eye movements to actions: Batsmen hit the ball. Nature Neuroscience, 3(12), 1340. https://doi.org/10.1038/81887

Lebeau, J.-C., Liu, S., Sáenz-Moncaleano, C., Sanduvete-Chaves, S., Chacón-Moscoso, S., Becker, B. J., & Tenenbaum, G. (2016). Quiet Eye and Performance in Sport: A Meta-Analysis. Journal of Sport and Exercise Psychology, 38(5), 441–457. https://doi.org/10.1123/jsep.2015-0123

Lee, D. N. (1998). Guiding movement by coupling taus. Ecological Psychology, 10(3–4), 221–250. https://doi.org/10.4324/9780203936672

Mann, D. L., Spratford, W., & Abernethy, B. (2013). The head tracks and gaze predicts: How the world’s best batters hit a ball. PLoS ONE, 8(3), e58289. https://doi.org/10.1371/journal.pone.0058289

Mann, D. Y., Williams, A. M., Ward, P., & Janelle, C. M. (2007). Perceptual-cognitive expertise in sport: A meta-analysis. Journal of Sport and Exercise Psychology, 29(4), 457–478. https://doi.org/10.1123/jsep.29.4.457

MLB Contracts. (2019). https://www.spotrac.com/mlb/contracts/sort-value/

Müller, S., & Fadde, P. J. (2016). The relationship between visual anticipation and baseball batting game statistics. Journal of Applied Sport Psychology, 28(1), 49–61. https://doi.org/10.1080/10413200.2015.1058867

Müller, S., Fadde, P. J., & Harbaugh, A. G. (2017). Adaptability of expert visual anticipation in baseball batting. Journal of Sports Sciences, 35(17), 1682–1690. https://doi.org/10.1080/02640414.2016.1230225

Muraskin, J., Sherwin, J., & Sajda, P. (2015). Knowing when not to swing: EEG evidence that enhanced perception-action coupling underlies baseball batter expertise. NeuroImage, 123, 1–10. https://doi.org/10.1016/j.neuroimage.2015.08.028

Murray, N., Kubitz, K., Roberts, C.-M., Hunfalvay, M., Bolte, T., & Tyagi, A. (2019). An examination of the oculomotor behavior metrics within a suite of digitized eye tracking tests. IEEE J Transl Eng Health Med., 1–5. https://righteye.com/wp-content/uploads/2019/02/Murray-Kubitz-Roberts-Hunfalvay-Bolte-Tyagi-2019.pdf

Paull, G., & Glencross, D. (1997). Expert perception and decision making in baseball. International Journal of Sport Psychology, 28, 35–56.

Poltavski, D., & Biberdorf, D. (2015). The role of visual perception measures used in sports vision programmes in predicting actual game performance in Division I collegiate hockey players. Journal of Sports Sciences, 33(6), 597–608. https://doi.org/10.1080/02640414.2014.951952

R Core Team. (2019). R: A language and environment for statistical computing. In R Foundation for Statistical Computing. https://doi.org/10.1017/CBO9781107415324.004

Regan, D. (1997). Visual factors in hitting and catching. Journal of Sports Sciences, 15(6), 533–558. https://doi.org/10.1080/026404197366985

RightEye. (2019). USA baseball and optometrists team up to improve players’ athletic abilities through sports vision. PR Newswire. https://www.prnewswire.com/news-releases/righteye-usa-baseball-and-optometrists-team-up-to-improve-players-athletic-abilities-through-sports-vision-300871734.html

Ripoll, H., & Fleurance, P. (1988). What does keeping one’s eye on the ball mean? Ergonomics, 31(11), 1647–1654. https://doi.org/10.1080/00140138808966814

Rubin, A., & Harris, W. F. (1995). Refractive variation during autorefraction: Multivariate distribution of refractive status. Optometry and Vision Science, 72(6), 403–410. https://doi.org/10.1097/00006324-199506000-00008

Rubin, D. B. (1987). Multiple imputation for nonresponse in surveys. John Wiley & Sons. https://doi.org/10.1002/9780470316696

Spering, M., & Gegenfurtner, K. R. (2008). Contextual effects on motion perception and smooth pursuit eye movements. Brain Research, 1225, 76–85. https://doi.org/10.1016/j.brainres.2008.04.061

Voss, M. W., Kramer, A. F., Basak, C., Prakash, R. S., & Roberts, B. (2010). Are expert athletes “expert” in the cognitive laboratory? A meta-analytic review of cognition and sport expertise. Applied Cognitive Psychology, 24(6), 812–826. https://doi.org/10.1002/acp.1588

Wang, L., Krasich, K., Bel-Bahar, T., Hughes, L., Mitroff, S. R., & Appelbaum, L. G. (2015). Mapping the structure of perceptual and visual-motor abilities in healthy young adults. Acta Psychologica, 157, 74–84. https://doi.org/10.1016/j.actpsy.2015.02.005

Watts, R. G., & Bahill, A. T. (1991). Keep your eye on the ball: Curve balls, knuckleballs, and fallacies of baseball. W. H. Freeman and Company.

Wilkins, L., & Appelbaum, L. G. (2019). An early review of stroboscopic visual training: Insights, challenges and accomplishments to guide future studies. International Review of Sport and Exercise Psychology, 1–16. https://doi.org/10.1080/1750984X.2019.1582081

Wood, S. N. (2004). Stable and efficient multiple smoothing parameter estimation for generalized additive models. Journal of the American Statistical Association, 99(467), 673–686. https://doi.org/10.1198/016214504000000980

