## Supplemental Contents for "Visual and oculomotor abilities predict professional baseball batting performance"

**Supplemental Content 1 (SC1)**

|  | **O-Swing Propensity (Model 3)** | |  | **Z-Swing Propensity (Model 2)** | |  | **League Level (Model 3)** | |
| --- | --- | --- | --- | --- | --- | --- | --- | --- |
|  | estimate(SE) | p value |  | estimate(SE) | p value |  | estimate(SE) | p value |
| Dynamic Visual Acuity (s) | -1.064(1.148) | 0.357 |  | -0.740(1.264) | 0.560 |  | -3.594(2.539) | 0.163 |
| Cardinal Reaction Time (s) | 1.503(3.305) | 0.651 |  | 2.939(3.531) | 0.409 |  | 6.147(6.392) | 0.341 |
| Simple Reaction Time (s) | 1.364(1.677) | 0.420 |  | 0.785(1.859) | 0.675 |  | -3.328(3.522) | 0.349 |
| Smooth Pursuit Accuracy (%) | -0.050(0.024) | 0.040* |  | -0.020(0.028) | 0.472 |  | 0.031(0.026) | 0.241 |
| General Oculomotor Latency (s) | 2.109(2.421) | 0.387 |  | 0.869(3.026) | 0.776 |  | -9.579(5.190) | 0.072 |
| General Oculomotor Speed (s) | 2.723(1.859) | 0.148 |  | 5.175(2.186) | 0.022* |  | -18.27(3.924) | < 0.001*** |
| General Processing Speed (s) | 4.199(2.059) | 0.046* |  | 6.043(2.706) | 0.030* |  | -5.861(4.069) | 0.156 |
| Visual Clarity (logMAR) |  |  |  | -2.280(1.293) | 0.084 |  |  |  |
| Contrast Sensitivity (log) |  |  |  | -0.049(0.656) | 0.940 |  |  |  |
| Near-Far Quickness (score) |  |  |  | 0.049(0.025) | 0.055 |  |  |  |
| Perception Span (score) |  |  |  | -0.006(0.012) | 0.597 |  |  |  |
| Multiple Object Tracking (score) |  |  |  | 0.000(0.000) | 0.558 |  |  |  |
| Reaction Time (s) |  |  |  | 10.19(6.078) | 0.099 |  |  |  |

*Slope parameter estimates (and SEs) and p values for the final regression models. Note: the Visual-Motor variables are on highly variable scales and therefore produce widely variable parameter estimates.*

**Supplemental Content 2 (SC2)**

|  | Model 1 | Model 2 | Model 3 | Null Model |
| --- | --- | --- | --- | --- |
| O-Swing Propensity | 32.86 (15.21, 50.82) | 32.63 (14.98, 50.63) | **27.92 (11.11, 46.25)** | 0 (0, .5.4) |
| Z-Swing Propensity | 22.52 (7.0, 41.16) | **21.56 (6.6, 39.81)** | 9.3 (0.55, 25.53) | 0 (0, .5.4) |
| Z-Miss Propensity | 21.22 (6.21, 39.71) | 18.72 (4.7, 37.06) | 3.05 (0.4, 15.86) | 0 (0, .5.4) |

*Model R^2^ estimates with 95% confidence intervals in parentheses (bolded if the associated models were chosen as final models).*

**Supplemental Content 3 (SC3)**

|  | *p* value | | |
| --- | --- | --- | --- |
|  | Model 1 vs. Model 2 | Model 2 vs. Model 3 | Model 1 vs. Model 3 |
| O-Swing Propensity | 0.74 | 0.78 | 0.9 |
| Z-Swing Propensity | 0.58 | 0.11 | 0.17 |
| Z-Miss Propensity | 0.23 | 0.07 | 0.08 |
| League Level | 0.1 | 0.64 | 0.43 |

*The p values from comparing vision models on a given outcome variable.*
